# Genome-wide Ultrabithorax binding analysis reveals highly targeted genomic loci at developmental regulators and a potential connection to Polycomb-mediated regulation

**DOI:** 10.1101/012609

**Authors:** Daria Shlyueva, Antonio C.A. Meireles-Filho, Michaela Pagani, Alexander Stark

## Abstract

Hox homeodomain transcription factors are key regulators of animal development. They specify the identity of segments along the anterior-posterior body axis in metazoans by controlling the expression of diverse downstream targets, including transcription factors and signaling pathway components. The *Drosophila melanogaster* Hox factor Ultrabithorax (Ubx) directs the development of thoracic and abdominal segments and appendages, and loss of Ubx function can lead for example to the transformation of third thoracic segment appendages (e.g. halters) into second thoracic segment appendages (e.g. wings), resulting in a characteristic four-wing phenotype. Here we present a *Drosophila melanogaster* strain with a V5-epitope tagged Ubx allele, which we employed to obtain a high quality genome-wide map of Ubx binding sites using ChIP-seq. We confirm the sensitivity of the V5 ChIP-seq by recovering 7/8 of well-studied Ubx-dependent cis-regulatory regions. Moreover, we show that Ubx binding is predictive of enhancer activity as suggested by comparison with a genome-scale resource of *in vivo* tested enhancer candidates. We observed densely clustered Ubx binding sites at 12 extended genomic loci that included ANTP-C, BX-C, Polycomb complex genes, and other regulators and the clustered binding sites were frequently active enhancers. Furthermore, Ubx binding was detected at known Polycomb response elements (PREs) and was associated with significant enrichments of Pc and Pho ChIP signals in contrast to binding sites of other developmental TFs. Together, our results show that Ubx targets developmental regulators via strongly clustered binding sites and allow us to hypothesize that regulation by Ubx might involve Polycomb group proteins to maintain specific regulatory states in cooperative or mutually exclusive fashion, an attractive model that combines two groups of proteins with prominent gene regulatory roles during animal development.

## INTRODUCTION

One of the most fascinating aspects of developmental gene regulation is the specification of animal body segment identity by homeobox domain containing transcription factors (TFs), including homeotic Hox factors. In many animals, Hox factors are arranged linearly in one or more genomic clusters and their sequential order along the genome sequence typically reflects their expression domain along the animals’ anterior-posterior axes. Their role in specifying segment identity has been revealed genetically by mutations in Hox factors that lead to homeotic transformations [1, 2]. For example, in *Drosophila melanogaster*, dominant mutations in the Antennapedia (Antp) locus lead to transformations of antennae to legs [3], recessive loss-of-function mutations in Antp transform the second leg into antenna [4], and Ultrabithorax (Ubx) mutations transform the balancing organs halteres into a second pair of wings [5, 6].

Such prominent phenotypes made the study of Hox factors and their regulatory targets important and attractive. Genetics established that Hox factors exhibit “posterior prevalence”, a regulatory hierarchy in which more posterior Hox genes repress more anterior ones and are dominant in specifying segment identity [2, 7]. A few direct targets and their regulatory elements have been identified and studied in detail [2, 8, 9] and microarray analyses after ubiquitous overexpression or misexpression of Hox factors have revealed putative regulatory targets genome-wide [10, 11]. Since extensive cross-regulation complicated the interpretation of gain- and loss-of-function studies, the binding site locations of Ubx and Dfd have been determined in *Drosophila* embryos and dissected imaginal discs by chromatin immunoprecipitation followed by microarray hybridization (ChIP-chip) [12-14] or by next-generation sequencing (ChIP-seq) [15]. These approaches were either based on antibodies against Ubx and Dfd [13-15] or made use of a protein trap line that contained a YFP insertion in the endogenous Ubx locus [12]. In this line, YFP appeared to recapitulate Ubx expression and flies homozygous or hemizygous for the Ubx-YFP allele were reported to exhibit reduced viability but only weak morphological phenotypes, which suggested that Ubx function was substantially normal [12].

The studies focused on the binding of Ubx in different tissues and/or analyzed the DNA sequence motifs, putative partner TFs, and chromatin features that are involved in the targeting of Ubx or Dfd to their binding sites [12, 13, 15]. The authors reported Ubx target gene networks, which for example confirmed that Ubx appeared to regulate several signaling pathways and dissected the *cis*-regulatory motif requirements and partner TFs involved in Ubx and Dfd binding and enhancer function.

Here we determine the location of Ubx binding sites in the entire genome of *D. melanogaster* embryos using ChIP-seq with antibodies against the heterologous V5 peptide and a *Drosophila melanogaster* strain in which we V5-epitope tagged the endogenous *Ultrabithorax* (*Ubx*) gene using homologous recombination. This revealed specific binding sites with high signal-to-noise ratios, which recovered 7 out of 8 known Ubx-dependent enhancers and were highly predictive of *in vivo* enhancer activity. Given the quality of the individual Ubx binding sites, we analyzed their genomic locations in detail, which suggests that the established regulation of other Hox genes by Ubx is direct and mediated via many individual Ubx binding sites. Ubx also binds in close proximity of many Polycomb complex genes and to known Polycomb response elements (PREs) and Ubx binding sites show significant enrichment of Polycomb and Pleiohomeotic binding genomewide, which we speculate could reflect a role of Hox genes in directing or antagonizing Polycomb-mediated developmental gene silencing.

## RESULTS

### Tagging of the endogenous Ubx locus by homologous recombination

To study Ubx binding throughout *Drosophila* embryogenesis, we first established a *Drosophila melanogaster* strain in which we tagged the endogenous *Ultrabithorax* (*Ubx*) gene with a V5 peptide using homologous recombination (Figure 1A). This strain, which is homozygous for the tagged Ubx allele (see below), should allow for ChIP with high sensitivity and specificity, independently of antibodies against the Ubx protein itself and without altering endogenous Ubx function. We chose to target the C-terminus that is shared between all known transcript isoforms and appears to allow the addition of peptide tags without impacting Ubx function [16].

**Figure 1.**
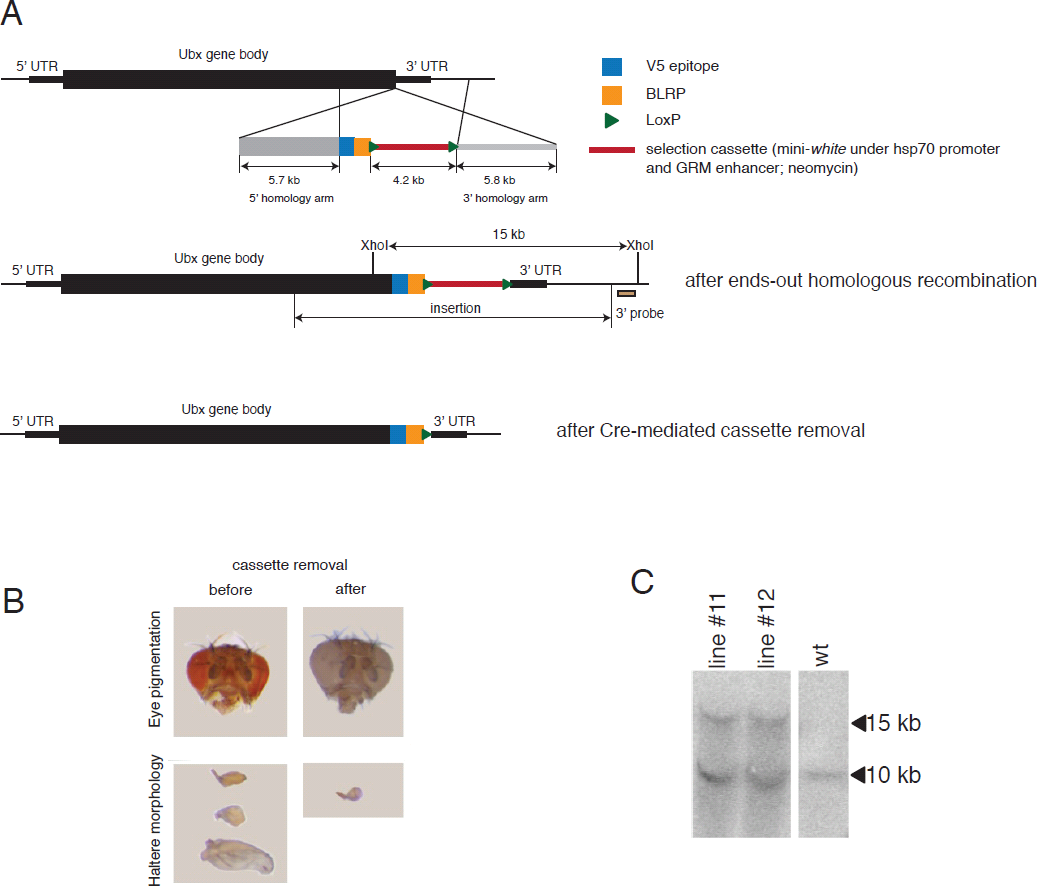
Creation of an epitope-tagged Ubx allele by homologous recombination. (A) Design of the targeting construct (top), the integration to the endogenous Ubx locus and the location of the probe for Southern blotting (middle) and the locus after the cassette removal (bottom). (B) The eye color and haltere morphology for candidate flies before (left) and after cassette removal (white-eyed fly) (right). (C) Southern blot confirming the correct integration for two independently recombined *Drosophila* lines #11 and #12 (heterozygous for the insert). w^1118^ flies were used as a control.

We first inserted the peptide tag and a selection cassette that was flanked by loxP sites using end-out homologous recombination [17-19] (Figure 1A and Materials and Methods). We selected flies that contained the targeting construct based on eye-color, which varied from dark red to orange, which might indicate various degrees of transcriptional repression of the selection marker in the flies’ eyes (Figure 1B). Interestingly, flies with orange eyes also had changes the morphology of their halteres, including increases in size and transformations to wings (Figure 1B), i.e. homeotic transformation characteristic for *Ubx* loss-of-function alleles [5, 6]. This suggested that the cassette was integrated correctly into the Ubx locus, which we confirmed by Southern blot analysis (Figure 1C). Importantly, the haltere phenotype was reversed when we removed the selection cassette (Figure 1B) and flies heterozygous or homozygous for the tagged allele both had wildtype haltere morphology, suggesting that the peptidetag – in contrast to the entire selection cassette – does not interfere with Ubx function. Taken together, we successfully tagged the 3’ end of *Ubx* and the tagged TF was functional as indicated by the wildtype phenotype in homozygous knockin flies.

### Characterization of genome-wide Ubx binding in Drosophila embryos

To determine Ubx binding sites genome-wide, we collected embryos of the homozygous tagged strain (0-16 hours post fertilization [hpf]) and performed ChIP-seq with an anti-V5 antibody. Two replicate ChIP-seq experiments from independent embryo collections showed strong and specific enrichments (peaks) and were highly similar with a genome-wide Pearson correlation coefficient [PCC] 0.86, demonstrating the reproducibility of the approach. We merged both replicates and identified genomic regions that were significantly enriched for Ubx binding (‘peaks’) with peakzilla [20]. We obtained 5282 peaks (peakzilla score ≥ 3), of which 1479 peaks were particularly strong with a score ≥ 5. To control for antibody-specificity and to obtain an estimate of the respective false-discovery rates for both score thresholds, we also performed the experiments with embryos from a non-tagged *D. melanogaster* strain (denoted hereafter as mock). This yielded 5 peaks with a score ≥ 3 and no peak with a score ≥ 5, demonstrating the specificity of the anti-V5 antibody and our approach and suggesting that the false discovery rates (FDRs) for peaks identified with the two score thresholds were 1.08% and 0.07%, respectively.

The Ubx binding sites were predominantly located in introns (41.8%) or intergenic regions (30.9%) and substantially depleted in coding regions and 3’UTRs, as expected for transcription factor binding sites and transcriptional enhancers [21-24] (Figure 2A). Importantly, the analysis recovered 7 out of 8 known Ubx-dependent *cis*-regulatory regions and binding sites near genes that loss- and gain-of-function studies suggested to be regulated by Ubx [10].

**Figure 2.**
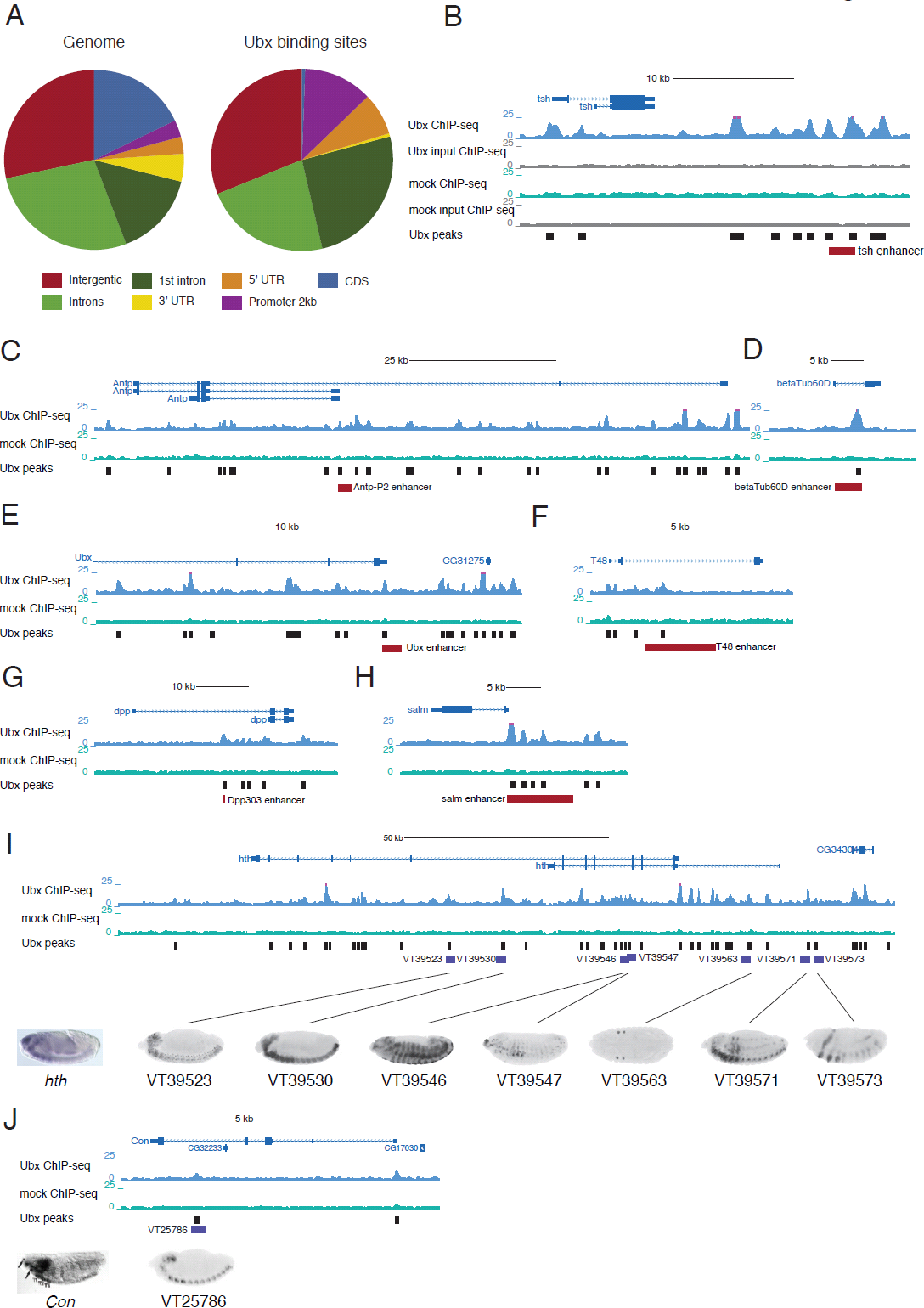
Genomic location of Ubx binding sites and recovery of known Ubx-dependent enhancers. (A) Genomic distribution of Ubx peaks (right) in comparison to the genome (left). (B-H) UCSC Genome Browser screenshots [75] of Ubx (blue), mock (green) ChIP-seq fragment density tracks and the Ubx peak calls at known Ubx-dependent enhancers (red bars) (see the main text for references). Panel (B) also contains the fragment density tracks for the two input samples (grey). (I) UCSC Genome Browser view of the *hth* locus and examples of Ubx-bound embryonic enhancers and their activity patterns [22] compared to *in situ* staining of the *hth* transcript [76]. (J) UCSC Genome Browser view of the *con* locus and Ubx-bound embryonic enhancer [22] compared to *in situ* staining of the *con* transcript [28].

The identified Ubx binding sites corroborate and provide putative molecular explanations for several long-standing observations, for example within Hox gene loci. Ubx binding to the promoter proximal part of *Antp*-P2 (Figure 2C) suggests that the proposed negative regulation by BX-C genes [25] could indeed be direct and mediated at least in part by Ubx. Similarly, binding of Ubx to its own promoter (Figure 1E) suggests that Ubx directly regulates its own expression, consistent with previous evidence that the *Ubx* promoter is involved in regulation of *Ubx* expression in the visceral mesoderm [26] and that this sequence can be bound by homeodomain-containing proteins [26, 27].

The first published chromatin immunopurification with anti-Ubx antibody revealed two transcripts directly regulated by Ubx: *Transcript 48* (*T48*) and 35 (or *connectin, con*) [28]. We confirmed that Ubx binds to the *T48* enhancer not only *in vitro* [29] but also *in vivo* (Figure 2F). In contrast, we did not observe binding to a putative *con* enhancer [30], and the respective DNA sequence indeed did not show any enhancer activity during embryogenesis [22]. Instead, we detected a Ubx binding site in a *con* intron and the corresponding sequence was active in the embryonic ventral nervous cord and in brain lobes, recapitulating *con* expression pattern in the nervous system [30] (Figure 2J). Similarly, while we did not detect binding at a putative *Dll* enhancer reported to be repressed by BX-C genes in abdominal segments [31], we observed Ubx binding sites more proximally to the *Dll* transcription starting site.

An intronic enhancer of *beta-tub60D* [32] was also bound by Ubx (Figure 2D), confirming the direct mode of regulation proposed previously based on *Ubx* gain- and loss-of-function experiments [32]. We also detected Ubx at well-characterized *tsh* (Figure 2A) and *dpp* enhancers (Figure 2G), which had been suggested to be positively regulated by Ubx based on DNaseI protection assays and enhancer assays of the wildtype enhancers and mutant variants [33, 34].

In addition to the small number of regulatory regions proposed to be under direct control of Ubx, hundreds of transcripts have been reported to respond to Ubx misexpression [10]. For example, *hth* was shown to be under negative control of Ubx and abd-A [35] and we indeed detected a large number of Ubx peaks in *hth* locus, many of which (17 out of 26) were active enhancers during embryogenesis with activity patterns reminiscent of *hth* expression [22] (Figure 2I).

Finally, several Ubx binding sites in a 10 kb embryonic enhancer upstream of *spalt major (salm)* [36] suggests that Ubx might regulate *salm* not only in haltere imaginal discs [37] but potentially already at embryonic stages (Figure 2H).

Taken together, we obtained high quality Ubx ChIP-seq data that confirmed previous observations regarding Hox-dependent gene regulation and provided further molecular insights into direct binding and regulation by Ubx .

### Ubx binds predominantly to active enhancers and Ubx binding is predictive of enhancer activity

The binding of TFs detected by ChIP-based methods does not imply functionality and not all TF-bound regions correspond to active enhancers [38, 39]. To test which proportion of Ubx binding sites coincide with active *cis*-regulatory elements, we used genome-scale resource of 7705 DNA fragments (Vienna tiles or VTs) tested in a reporter assay and imaged throughout *Drosophila* embryogenesis [22].

77% of all VTs that contained at least one Ubx peak summit (score ≥ 5) were active, a 1.7-fold increase compared to all VTs of which 46% were active [22] (Figure 3A). The increase was even more prominent when examining the fraction of active VTs at each of the developmental stage intervals separately, which was on average 2.5 times higher for Ubx-bound VTs (Figure 3A).

**Figure 3.**
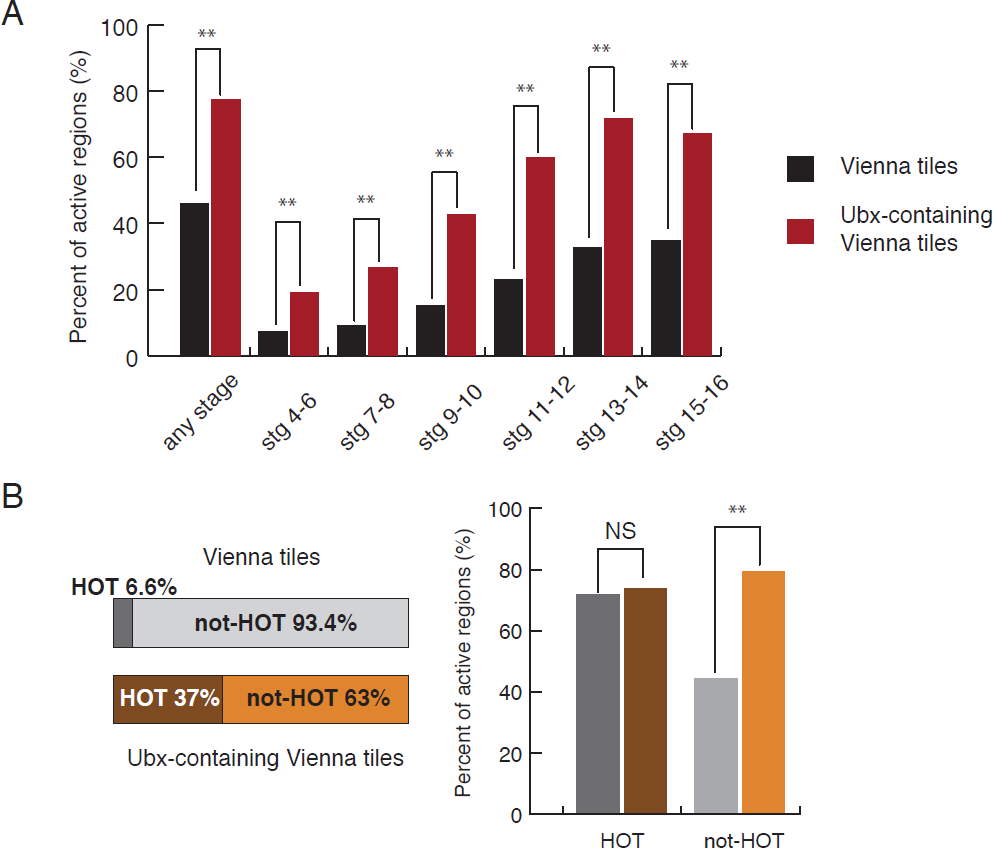
Ubx binds to active enhancers. (A) The bar plot shows the percentage of Vienna tiles (VTs) without or with Ubx binding sites (black and red bars, respectively) that is active at any stage of embryogenesis (left) or at the indicated embryonic stages. Hypergeometric p-value: **P<10^−10^. (B) The left panel shows the fraction of all VTs (top) and the fraction of Ubx-bound VTs (bottom) that overlaps HOT regions (dark shading). The bar plot on the right shows the percentage of active tiles for the four subsets of VTs defined on the left panel. NS – not significant. Hypergeometric p-value: **P<10^−19^.

It was recently shown that TF binding detected by ChIP had a tendency to accumulate at specific genomic regions, termed HOT regions (highly occupied targets) [14, 38]. Interestingly, such regions were shown to function as transcriptional enhancers in *Drosophila*, but the functional contribution of each bound TF remained unclear as the HOT regions’ activity patterns did not always coincide or were consistent with the bound TFs’ expression patterns [38]. Our observation that Ubx binding was predictive of enhancer activity was also true when we analyzed the 63% VTs that contained Ubx binding sites but no HOT regions separately (Figure 3B): 79% of Ubx-bound VTs that did not contain any HOT region were active compared to only 44% of all such VTs. The difference was much less pronounced for Ubx-bound VTs that also contained HOT regions (Figure 3B), as HOT regions are frequently active more generally [38].

### Multiple Ubx binding sites in Hox gene loci

One of the prominent features of Hox factors is their extensive cross-regulation [7, 8, 40]. For example, Ubx was shown to regulate its own transcription [26] and that of *Antp* [25]. We therefore first analyzed Ubx binding sites within the ANTP-C and BX-C loci. Interestingly, the 350kb ANTP-C between *lab* and *Antp* contained 50 Ubx binding sites (score ≥ 3), of which 22 were strongly bound (score ≥ 5), a substantial enrichment compared to the 3 binding sites we observed per 100kb window on average (4.8-fold; Poisson P-value P=1.8 × 10^−18^). Furthermore, the 340kb BX-C locus between *Ubx* and *abd-B* contained 73 binding sites, 43 of which were strong (Figure 4A). Importantly, only 30% and 19% of the Ubx binding sites in ANTP-C and BX-C, respectively, coincided with highly occupied target (HOT) regions [14, 38], suggesting that the observed enrichment was specific to Ubx and neither due to cross-linking artifacts [41-43] nor shared by many other TFs.

**Figure 4.**
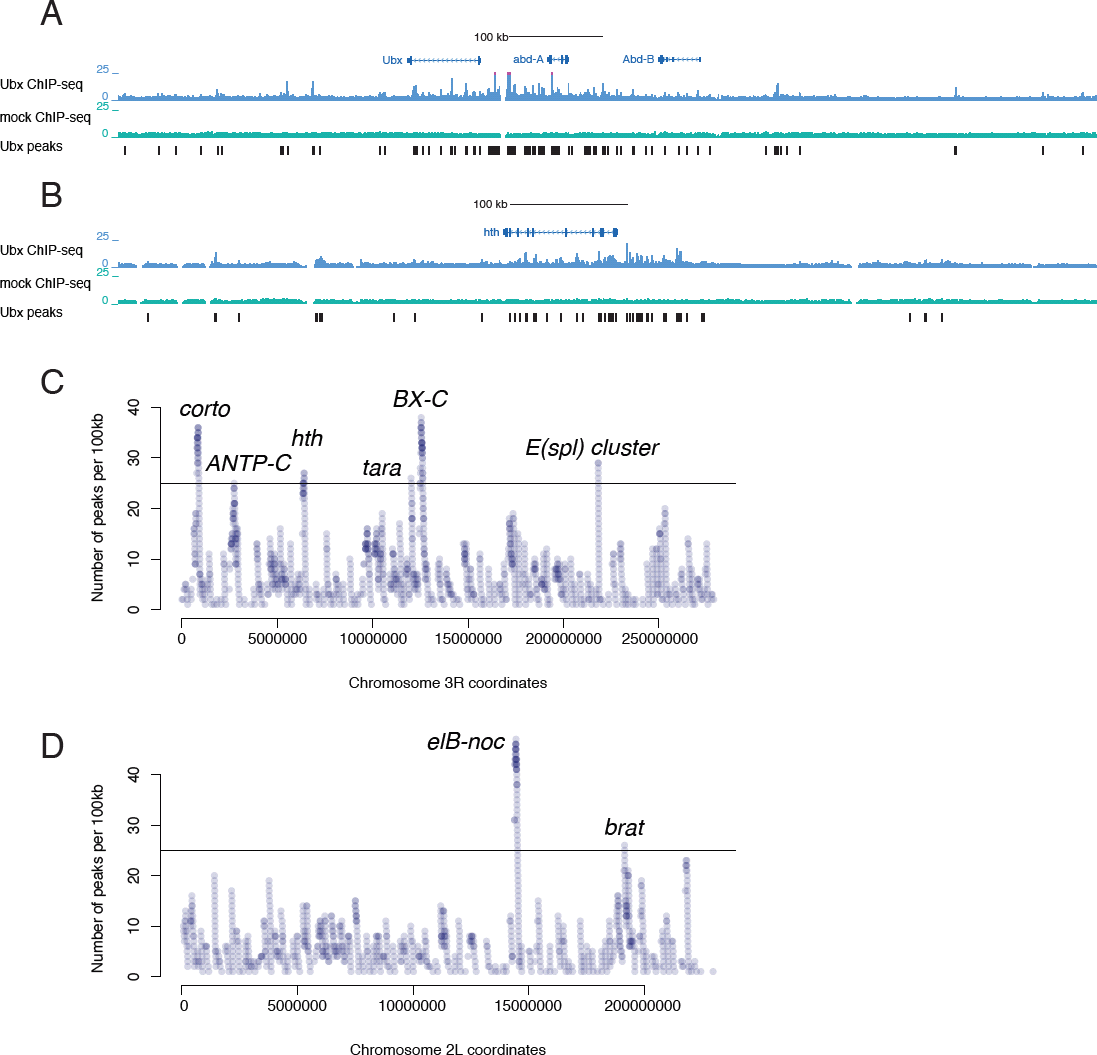
Clustered Ubx binding sites at the genomic loci of important developmental regulators. (A, B) UCSC Genome Browser screenshots at BX-C and *hth* gene loci show ChIP-seq fragment density tracks and Ubx peak calls, revealing many clustered binding sites. (C, D) The number of Ubx peaks per 100 kb on chromosomes 3R and 2L (each 100 kb window starts at a Ubx peak, covering all possible windows that contain at least one Ubx peak). The plots for chromosomes 2R, 3L, X and 4 are in Figure S1. Representative genes for all windows containing ≥25 Ubx peaks are labeled.

### Clustered Ubx binding sites at highly targeted genomic loci (HTGLs) around developmental regulators and genes related to the Polycomb complex (Pc)

To systematically determine genomic regions that contain clustered Ubx binding sites, we counted the number of peaks in 100 kb windows genome-wide (Figure 4C, D and Figure S1). This revealed two non-overlapping 100 kb windows with 25 Ubx binding sites (score ≥ 3) or more on chromosome 2L, one on chromosome 2R, and three and six on chromosomes 3L and 3R, respectively (Figure 4C, D and Figure S1).

The two windows with prominent Ubx binding site clusters on chromosome 2L overlapped the gene loci of *elB-noc* and *brat* (Figure 4D). *elB* and *noc* were suggested to play a role in cell proliferation [44] and necessary for the appendage formation [45]. *Brat* is known to regulate post-transcriptional gene expression [46, 47] and its mutations caused defects in abdominal segments [47]. The Ubx binding site cluster on chromosome 2R spanned the *sbb* and *tango8* locus, and the clusters on 3L are near the apoptotic genes *scyl* and *chrb* and *W, grim* and *rpr. Scyl* and *chrb* were previously shown to be de-repressed in *Ubx, abd-A* and *Abd-B* mutant flies [48] (Figure S1). Apoptosis is necessary for the maintenance of segments boundaries, and Ubx – similarly to Dfd and Abd-B [49] – might be linked to its regulation. The third Ubx-rich cluster on chromosome 3L contained *tonalli* (*tna*), a Trithorax group gene that was identified together with *taranis (tara)* (see below) and mutations of which induced homeotic transformations [50]. Besides clustered Ubx binding sites in BX-C, chromosome 3R contained multiple Ubx binding at the *hth* locus (Figure 4B). Hth is a known partner of Hox factors, which has been reported to modulate the specificity of Hox factor binding *in vivo* [9], and our data suggest that Ubx might directly regulate *hth* via a large number of binding sites. Another noticeable cluster on 3R is in the *Enhancer of split (E(spl))* complex, which is a genomic cluster of basic helix-loop-helix (bHLH) transcription factors that are involved in Notch signaling and which are regulated by Ubx in haltere [12, 51](Figure 4C).

Chromosome 3R also contained two Ubx binding site clusters with 36 and 26 binding sites per 100kb near the *corto* and *taranis* (*tara*) gene loci (Figure 4C). Corto and Tara are Polycomb- and Trithorax-interacting proteins, respectively [52-54] and mutant alleles of both genes were shown to enhance the Polycomb/Thritorax mutant phenotypes and affect Hox gene regulation [54, 55]. This is particularly interesting, as we observed multiple Ubx binding sites also in the gene loci of many Polycomb and Trithorax complex members (e.g. *trx, osa, ash2, Pc, Pcs, ph-p* etc.; Figure 5D). In addition, several of the gene loci bound by Ubx are known direct targets of the Polycomb complex, including *elB-noc* locus [56] and *sbb/tango8*, which contain a predicted PRE element [57] (see also below).

**Figure 5.**
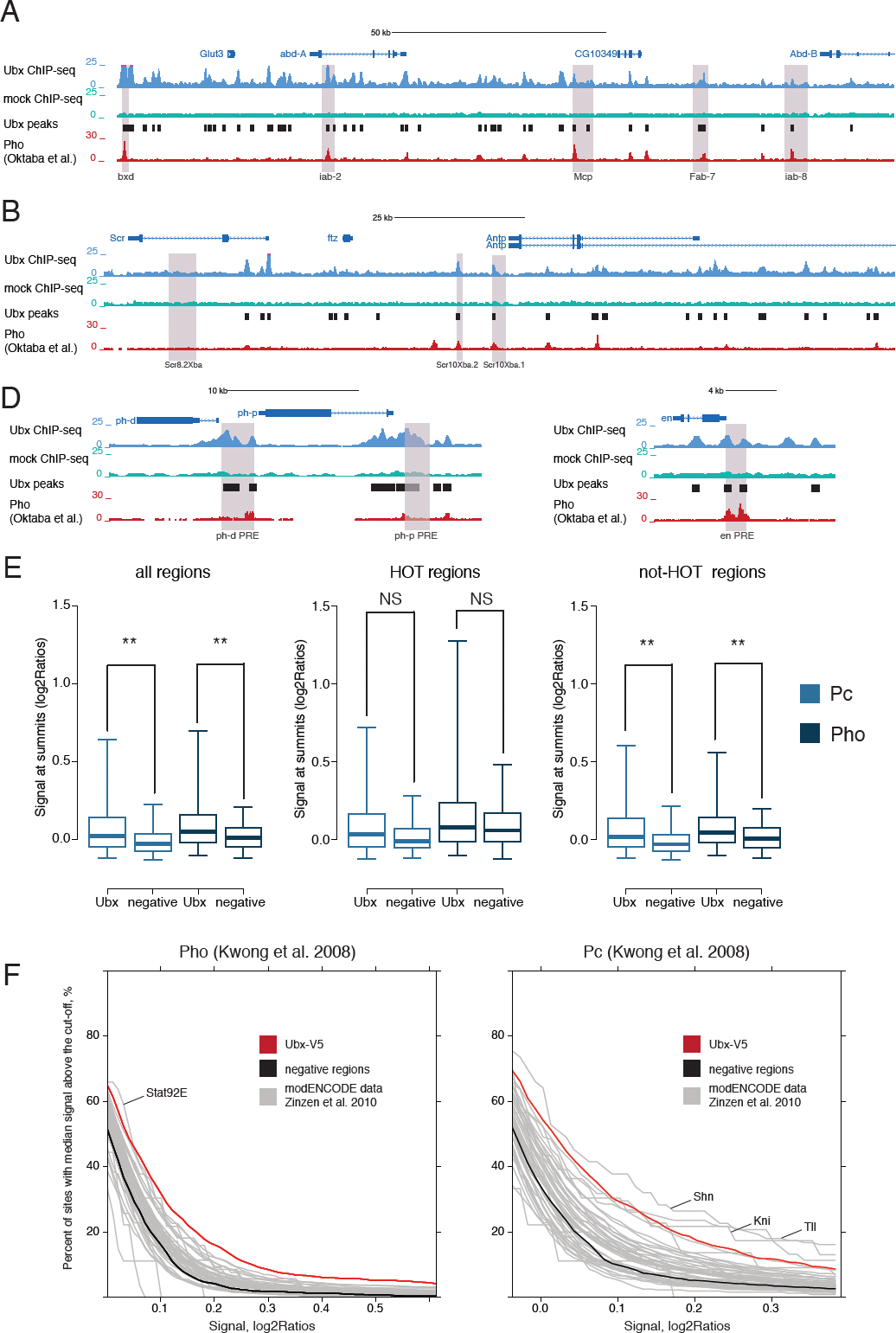
Strong overlap of Ubx and Polycomb complex binding sites in entire embryos. (A-D) UCSC Genome Browser screenshots of Ubx and mock ChIP-seq tracks at known PRE/TRE elements (purple shading) at BX-C, ANTP-C, *en, ph-p* and *ph-d.* The coordinates are from [57]. Pho track is from [65]. (E) The box plots show the Pc and Pho ChIP-chip signal (ChIP/input ratio [log2]) [64] at the Ubx peak summits and at control regions. The left panel represents all regions, the middle panel positions that overlap HOT regions and the right panel those that do not overlap HOT regions. NS – not significant; Wicoxon test: **P<10^−20^; equivalent plots for other TFs are in Figure S3. (F) The plots show the percentage of TFs binding sites that have Pc or Pho signal [64] greater than a given threshold value (X-axis; red line: Ubx, black: control regions and grey: other TFs from [14, 24]).

The occurrence of such highly targeted genomic loci (HTGLs) and their coincidence with important developmental regulators is striking. To assess the functional relevance of the Ubx binding sites in HTGLs, we evaluated their enhancer activities in transgenic embryos [22] and compared them to the activities of Ubx-bound regions outside HTGLs. Interestingly, VTs overlapping at least one of the Ubx-binding sites within HTGLs were significantly more often active during embryogenesis than those overlapping Ubx binding sites outside HTGLs (64 out of 78 [82%] vs 177 out of 264 [67%], *P*-value=0.0066).

Our observation that Ubx binds to clustered sites within HTGLs around important developmental regulators and that most of these binding sites correspond to active enhancers is interesting. While multiple closely adjacent binding sites of homeodomain proteins within a single enhancer might assist cooperative interactions between TFs and assure stable interaction with the enhancer DNA [58], the HTGLs reported here correspond to clusters of several individual enhancers, presumably similar to locus control regions (LCRs; [59, 60]) or super enhancers [61]. The control of developmental regulators via many densely spaced enhancers that are bound by Ubx is also interesting given that Ubx itself is an important developmental regulator that determines segment identity.

### A putative link between Ubx and Polycomb targeting

Hox factors are among the best-characterized targets of the Polycomb and Trithorax complexes, which function to maintain repressive or active transcriptional states, respectively throughout development [62, 63]. In *Drosophila*, they have been reported to act through specialized genomic elements, called Polycomb or Trithorax response elements (PRE/TREs; [62]), and BX-C contains several well-studied PREs [62, 63].

One of the Ubx peaks with highest ChIP enrichment genome-wide co-localized with a known PRE/TRE (genomic coordinates from [57]) near the non-coding gene *bxd* (Figure 5A). Moreover, other well-characterized PREs in Hox loci and near *ph-d, ph-p* and *en* [57] also all contained Ubx peaks (Figure 5A-D).

Given the small number of *in vivo* validated PRE/TREs, we used genome-wide binding data for Polycomb (Pc) and Pleiohomeotic (Pho) [64] to assess more systematically whether Ubx-bound regions were associated with Polycomb complexes genome-wide. Indeed, the enrichment of Pc and Pho binding was higher at Ubx peaks than at control regions (P<10^−29^) (Figure 5E, left panel). The same was true for Ubx binding sites outside HOT regions (P<10^−20^) (Figure 5E, right panel), suggesting that observed association with Pc and Pho was specific to Ubx and did not result from general accessibility of the binding sites. Indeed, if we assess Pc and Pho ChIP enrichments at binding sites for Ubx and 43 different developmental TFs [14, 24], they were higher on average at Ubx binding sites (Figure S3) and the proportion of TF binding sites with Pc and Pho ChIP signals greater than a specific cut-off was higher for Ubx across a wide range of cut-offs (Figure 5F and S2). The same was true when analyzing independent dataset for Polycomb-associated proteins binding and the Polycomb-associated histone modification H3K27me3 (Figure S2).

Considering the duality of PRE/TREs, which can switch from activation to repression and our observation that Ubx binding is predictive of active enhancers, we decided to assess the ability of PRE/TREs to enhance expression of a reporter depending on the presence of Ubx. For this we evaluated the activities of VT fragments overlapping 407 PREs (as defined by Pho binding in embryos [65]). Interestingly, 63.4% of Pho-bound VTs (71 out of 112) were active at any stage of embryonic development in comparison to the 46% positive rate for VTs overall (hypergeometric *P*-value=1.4 × 10^−8^). The percentage of regions acting as enhancers in *Drosophila* embryos increased to 72.0% (31 out of 43) when considering VTs co-bound by Pho and Ubx in contrast to 58.6% (41 out of 70) for VTs bound only by Pho. It suggests that Ubx acts jointly or mutually exclusively with Pc proteins on putative PRE/TRE leading to activation of such genomic regions.

Our ChIP-seq data from entire embryos and different embryonic stages show that Ubx and the Polycomb complex bind to the same genomic regions, suggesting a dynamic interplay between Ubx and Polycomb recruitment. This could occur in parallel or spatially or temporally exclusive domains with different mechanistic implications as we discuss below.

## DISCUSSION

Here we present a *Drosophila melanogaster* strain with a Ubx allele that is tagged at the Ubx C-terminus with a V5 peptide and allowed us to study Ubx binding genome-wide. The tag also contains a biotin-ligase-recognition peptide (BLRP), which should be useful for biochemical approaches, including the biotin/streptavidin-based purification of Ubx containing protein complexes [66, 67] and might allow – combined with the targeted expression of biotin ligase (BirA) – to perform tissue-specific ChIP-seq experiments.

We further present a high-quality ChIP-seq dataset that allowed the identification of individual Ubx binding sites genome-wide. These binding sites frequently overlap with active enhancers and Ubx binding is predictive of enhancer activity, especially outside HOT regions. Importantly, Ubx binds extensively to HTGLs, which often overlap the gene loci of developmental regulators and genes that are regulated by the Polycomb complex and the majority of these binding sites are functioning as enhancers during embryogenesis.

Our observation that Ubx binds to known PREs/TREs and that Ubx binding sites also show a significant Pc and Pho ChIP signal is suggestive of a model in which Ubx could be upstream of Pc targeting and involved in mediating or antagonizing Pc and Pho recruitment to their genomic binding sites. The data are consistent with two scenarios: Ubx and Pc/Trx binding might occur predominantly in the same cells and Ubx could be involved in recruiting Pc/Trx to their binding sites. Alternatively, Ubx and Pc/Trx might occur predominantly in mutually exclusive spatial domains or at different stages in the developing embryo and Ubx could potentially counteract Pc binding.

The first hypothesis is consistent with known Polycomb-dependent Ubx repression by high transient levels of Ubx in haltere [68] and the known repression of *bxd* in Ubx-expressing cells, which involved components of the Trx complex [69]. Our finding that Ubx was bound at *bxd* locus suggests that this repression could be direct and mediated by the Hox factor.

On the other hand, Ubx binding has not been observed at the *Abd-A* and *abd-B* loci in haltere [12], a tissue in which *Abd-A* and *abd-B* are repressed by Pc. Similarly, sites that are bound by Ubx in embryos have high levels of Pc and H3K27me3 in S2 cells (Figure S2) that do not express Ubx [70]. Therefore, while Ubx could be involved transiently during initial steps of Pc recruitment, it does not seem to be required for repression and Pc might even restrict TF access to these loci [12]. Moreover, as Pc is typically associated with repression, the strong enrichment of active enhancers at Ubx binding sites suggests that Ubx could counteract Pc, potentially through enhancer activation. In other cells, Pc would then bind to and silence the same regions thereby counteracting Ubx function, leading for example to the high levels of H3K27me3 observed in ChIP experiments from entire embryos.

The prediction that Ubx might be involved in specifying or counteracting the recruitment of Polycomb to specific genomic loci is attractive as it links Hox genes, which are involved in the definition of segment identity with Polycomb, which has been implicated in the maintenance of transcriptional regulatory states throughout development. While we find that several TFs co-localize with Pc/Pho binding sites in ChIP from entire embryos, Ubx had the most prominent effect. Given the attractiveness and potential importance of this link between Hox genes and Polycomb, we would like to share this observation with the broader scientific community.

### Materials and Methods

#### Drosophila stocks

Drosophila flies were kept at 25° C on standard food. w^1118^ strain (denoted as mock) was obtained from the Bloomington Stock center.

#### Generation of the donor constructs and homologous recombination

The detailed description of P[acman] vector modifications are in {MeirelesFilho:2013ka}. The genomic coordinates of 5’ homology arm: chr3R 12484497 – 12490226 ; 3’ homology arm: chr3R 12478739 - 12484493. The tag included the V5 epitope (see below; in bold), biotin-ligase-recognition-peptide (BLRP) (see below; in italic) separated by PreScission cleavage site (see below; underscored). The DNA sequence of the tag: GCGGCGGCG**GCAAGCCCATCCCCAACC**<u>CCCTGCTGGGCCTGGATAGCACCCTG</u>GAGGTGCTGTTCCAGGGCCCCGAGAACCTGTACTTCCAGGGC*ATGGCCAGCAGCCTGCGCCAG ATCCTGGATAGCCAGAAGATGGAGTGGCGCAGCAACGCCGGCGGCAGC*tgaGGTACC. The selection cassette consisted of mini-white gene under hsp70 promoter and GRM enhancer and 2 flanking LoxP sites. The construct was injected into ZH-attP-51D fly strain with the landing site on chr2R [71]. Genetic crosses were done as described in [19]. Positive candidates were confirmed by Southern blot: the restriction enzyme used: XhoI (NEB); 3’ end probe: chr3R 12477423 -12478376). The selection cassette removal was done by Cre-mediated recombination [19].

#### Embryo collection and ChIP-seq

The Ubx-tagged and w^1118^ flies were kept in large populational cages at 25°C. Embryos were collected for 16 hrs (overnight), dechorionated and frozen. Approximately 1g of frozen embryos was fixed and processed as described [72]. Nuclei were sonicated in 1.5 ml of nuclear lysis buffer [73] with the Tip sonicator (Omni Sonic Ruptor 250 Watt Ultrasonic Homogenizer) for 7 cycles (1 min on [Duty cycle 30%, Output 20%], 1 min off). The average size of sheared fragments was approximately 500 bp. 500 μl of sonicated chromatin was incubated with 25 μl of blocked anti-V5 agarose affinity gel (Sigma, A7345-1ML) and 500 μl of RIPA buffer [73] for 2 hrs at 4°C. The beads were washed as described [73]. A total of 3 ng of material was used for library generation.

#### In vivo enhancer activity analysis

All enhancer activity assays are based on transcriptional reporter assays in transgenic embryos available from the Vienna Tile (VT) resource [22] at http://enhancers.starklab.org.

#### Deep-sequencing

Sequencing was performed at the CSF NGS Unit (http://www.csf.ac.at) on an Illumina HiSeq2000 machine. We processed data as single-end sequencing data, compared two independent biological replicates and merged them for the subsequent analyses.

#### Reads processing and peak calling

We obtained unique fragments by mapping reads to dm3 genome using bowtie [74], allowing maximum three mismatches. Significantly enriched regions (peaks) were identified using peakzilla [20] with default settings. As a cut-off parameter we used a peak score that takes into account the enrichment and distribution of reads in a peak region [20].

## Figure legends

**Figure S1. Clustered Ubx binding sites at the loci of important developmental genes.** The number of Ubx peaks per 100 kb on chromosomes 2R, 3L, X and 4 (as in main Figure 4C,D). Each 100 kb window starts at a Ubx peak, covering all possible windows that contain at least one Ubx peak. Representative genes for windows containing ≥25 Ubx peaks are indicated.

**Figure S2. Association of Polycomb complex with binding sites for developmental transcription factors (TFs).** The plots show the percentage of TF binding sites for which the ChIP signal for the indicated Polycomb group protein or the Polycomb-associated histone modification H3K27me3 [65, 70, 77] is greater than a given threshold value (X-axis; as in main Figure 5F).

**Figure S3. Strong association of Pc and Pho with Ubx binding sites.** The boxplots show the distributions of Pc and Pho normalized ChIP signal (ChIP/input ratio [log2]) at the binding sites of the indicated transcriptional factors (as in Figure 5E). The binding sites are from [14] and [24] and are restricted to TFs that had Pc and Pho signals significantly higher than control regions. NS – not significant; Wilcoxon test: \**P*<0.001, \*\**P*<10^−5^; #: TF binding sites from [24].

## References

1. Lewis EB: A gene complex controlling segmentation in Drosophila. Nature 1978, 276:565–570.

2. Pearson JC, Lemons D, McGinnis W: Modulating Hox gene functions during animal body patterning. Nat Rev Genet 2005, 6:893–904.

3. Kaufman TC, Lewis R, Wakimoto B: Cytogenetic Analysis of Chromosome-3 in Drosophila-Melanogaster - the Homoeotic Gene-Complex in Polytene Chromosome Interval 84a-B. Genetics 1980, 94:115–133.

4. Struhl G: A homoeotic mutation transforming leg to antenna in Drosophila. 1981.

5. Struhl G: Genes controlling segmental specification in the Drosophila thorax. Proc Natl Acad Sci USA 1982, 79:7380–7384.

6. Weatherbee SD, Halder G, Kim J, Hudson A, Carroll S: Ultrabithorax regulates genes at several levels of the wing-patterning hierarchy to shape the development of the Drosophila haltere. Genes Dev 1998, 12:1474–1482.

7. McGinnis W, Krumlauf R: Homeobox genes and axial patterning. Cell 1992, 68:283–302.

8. Hueber SD, Lohmann I: Shaping segments: Hox gene function in the genomic age. Bioessays 2008, 30:965–979.

9. Mann RS, Lelli KM, Joshi R: Hox specificity unique roles for cofactors and collaborators. Curr Top Dev Biol 2009, 88:63–101.

10. Hueber SD, Bezdan D, Henz SR, Blank M, Wu H, Lohmann I: Comparative analysis of Hox downstream genes in Drosophila. Development 2007, 134:381–392.

11. Pavlopoulos A, Akam M: Hox gene Ultrabithorax regulates distinct sets of target genes at successive stages of Drosophila haltere morphogenesis. Proc Natl Acad Sci USA 2011, 108:2855–2860.

12. Choo SW, White R, Russell S: Genome-wide analysis of the binding of the Hox protein Ultrabithorax and the Hox cofactor Homothorax in Drosophila. PLoS ONE 2011, 6:e14778.

13. Slattery M, Ma L, Négre N, White KP, Mann RS: Genome-wide tissue-specific occupancy of the hox protein ultrabithorax and hox cofactor homothorax in Drosophila. PLoS ONE 2011, 6:e14686.

14. modENCODE Consortium, Roy S, Ernst J, Kharchenko PV, Kheradpour P, Nègre N, Eaton ML, Landolin JM, Bristow CA, Ma L, Lin MF, Washietl S, Arshinoff BI, Ay F, Meyer PE, Robine N, Washington NL, Di Stefano L, Berezikov E, Brown CD, Candeias R, Carlson JW, Carr A, Jungreis I, Marbach D, Sealfon R, Tolstorukov MY, Will S, Alekseyenko AA, Artieri C, et al.: Identification of functional elements and regulatory circuits by Drosophila modENCODE. Science 2010, 330:1787–1797.

15. Sorge S, Ha N, Polychronidou M, Friedrich J, Bezdan D, Kaspar P, Schaefer MH, Ossowski S, Henz SR, Mundorf J, Rätzer J, Papagiannouli F, Lohmann I: The cis-regulatory code of Hox function in Drosophila. EMBO J 2012.

16. Ronshaugen M, McGinnis N, McGinnis W: Hox protein mutation and macroevolution of the insect body plan. Nature 2002, 415:914–917.

17. Gong WJ, Golic KG: Ends-out, or replacement, gene targeting in Drosophila. Proc Natl Acad Sci USA 2003, 100:2556–2561.

18. Venken KJT, He Y, Hoskins RA, Bellen HJ: P[acman]: a BAC transgenic platform for targeted insertion of large DNA fragments in D. melanogaster. Science 2006, 314:1747–1751.

19. Huang J, Zhou W, Watson AM, Jan Y-N, Hong Y: Efficient Ends-Out Gene Targeting In Drosophila. Genetics 2008, 180:703–707.

20. Bardet AF, Steinmann J, Bafna S, Knoblich JA, Zeitlinger J, Stark A: Identification of transcription factor binding sites from ChIP-seq data at high resolution. Bioinformatics 2013, 29:2705–2713.

21. Zeitlinger J, Zinzen RP, Stark A, Kellis M, Zhang H, Young RA, Levine M: Whole-genome ChIP-chip analysis of Dorsal, Twist, and Snail suggests integration of diverse patterning processes in the Drosophila embryo. Genes Dev 2007, 21:385–390.

22. Kvon EZ, Kazmar T, Stampfel G, Yáñnez-Cuna JO, Pagani M, Schernhuber K, Dickson BJ, Stark A: Genome-scale functional characterization of Drosophila developmental enhancers in vivo. Nature 2014.

23. Arnold CD, Gerlach D, Stelzer C, Boryn LM, Rath M, Stark A: Genome-Wide Quantitative Enhancer Activity Maps Identified by STARR-seq. Science 2013.

24. Zinzen RP, Girardot C, Gagneur J, Braun M, Furlong EEM: Combinatorial binding predicts spatio-temporal cis-regulatory activity. Nature 2009, 462:65–70.

25. Boulet AM, Scott MP: Control elements of the P2 promoter of the Antennapedia gene. Genes Dev 1988, 2:1600–1614.

26. Müller J, Thüringer F, Biggin M, Züst B, Bienz M: Coordinate action of a proximal homeoprotein binding site and a distal sequence confers the Ultrabithorax expression pattern in the visceral mesoderm. EMBO J 1989, 8:4143–4151.

27. Bienz M, Saari G, Tremml G, Müller J, Züst B, Lawrence PA: Differential regulation of Ultrabithorax in two germ layers of Drosophila. Cell 1988, 53:567–576.

28. Gould AP, Brookman JJ, Strutt DI, White RA: Targets of homeotic gene control in Drosophila. Nature 1990, 348:308–312.

29. Strutt DI, White RA: Characterisation of T48, a target of homeotic gene regulation in Drosophila embryogenesis. Mech Dev 1994, 46:27–39.

30. Gould AP, White RA: Connectin, a target of homeotic gene control in Drosophila. Development 1992, 116:1163–1174.

31. Gebelein B, McKay DJ, Mann RS: Direct integration of Hox and segmentation gene inputs during Drosophila development. Nature 2004, 431:653–659.

32. Hinz U, Wolk A, Renkawitz-Pohl R: Ultrabithorax is a regulator of beta 3 tubulin expression in the Drosophila visceral mesoderm. Development 1992, 116:543–554.

33. McCormick A, Coré N, Kerridge S, Scott MP: Homeotic response elements are tightly linked to tissue-specific elements in a transcriptional enhancer of the teashirt gene. Development 1995, 121:2799–2812.

34. Capovilla M, Brandt M, Botas J: Direct regulation of decapentaplegic by Ultrabithorax and its role in Drosophila midgut morphogenesis. Cell 1994, 76:461–475.

35. Kurant E, Pai CY, Sharf R, Halachmi N, Sun YH, Salzberg A: Dorsotonals/homothorax, the Drosophila homologue of meis1, interacts with extradenticle in patterning of the embryonic PNS. Development 1998, 125:1037–1048.

36. Kühnlein RP, Brönner G, Taubert H, Schuh R: Regulation of Drosophila spalt gene expression. Mech Dev 1997, 66:107–118.

37. Galant R, Walsh CM, Carroll SB: Hox repression of a target gene: extradenticle-independent, additive action through multiple monomer binding sites. Development 2002, 129:3115–3126.

38. Kvon EZ, Stampfel G, Yáñez-Cuna JO, Dickson BJ, Stark A: HOT regions function as patterned developmental enhancers and have a distinct cis-regulatory signature. Genes Dev 2012, 26:908–913.

39. Li X-Y, MacArthur S, Bourgon R, Nix D, Pollard DA, Iyer VN, Hechmer A, Simirenko L, Stapleton M, Luengo Hendriks CL, Chu HC, Ogawa N, Inwood W, Sementchenko V, Beaton A, Weiszmann R, Celniker SE, Knowles DW, Gingeras T, Speed TP, Eisen MB, Biggin MD: Transcription factors bind thousands of active and inactive regions in the Drosophila blastoderm. PLoS Biol 2008, 6:e27.

40. Hafen E, Levine M, Gehring WJ: Regulation of Antennapedia transcript distribution by the bithorax complex in Drosophila. Nature 1984, 307:287–289.

41. Teytelman L, Thurtle DM, Rine J, van Oudenaarden A: Highly expressed loci are vulnerable to misleading ChIP localization of multiple unrelated proteins. Proc Natl Acad Sci USA 2013, 110:18602–18607.

42. Park D, Lee Y, Bhupindersingh G, Iyer VR: Widespread misinterpretable ChIP-seq bias in yeast. PLoS ONE 2013, 8:e83506.

43. Poorey K, Viswanathan R, Carver MN, Karpova TS, Cirimotich SM, McNally JG, Bekiranov S, Auble DT: Measuring chromatin interaction dynamics on the second time scale at single-copy genes. Science 2013, 342:369–372.

44. Tseng A-SK, Hariharan IK: An overexpression screen in Drosophila for genes that restrict growth or cell-cycle progression in the developing eye. Genetics 2002, 162:229–243.

45. Weihe U: Proximodistal subdivision of Drosophila legs and wings: the elbow-no ocelli gene complex. Development 2004, 131:767–774.

46. Betschinger J, Mechtler K, Knoblich JA: Asymmetric Segregation of the Tumor Suppressor Brat Regulates Self-Renewal in Drosophila Neural Stem Cells. Cell 2006, 124:1241–1253.

47. Sonoda J, Wharton RP: Drosophila Brain Tumor is a translational repressor. Genes Dev 2001, 15:762–773.

48. Scuderi A, Simin K, Kazuko SG, Metherall JE, Letsou A: scylla and charybde, homologues of the human apoptotic gene RTP801, are required for head involution in Drosophila. Dev Biol 2006, 291:110–122.

49. Lohmann I, McGinnis N, Bodmer M, McGinnis W: The Drosophila Hox gene deformed sculpts head morphology via direct regulation of the apoptosis activator reaper. Cell 2002, 110:457–466.

50. Gutiérrez L, Zurita M, Kennison JA, Vázquez M: The Drosophila trithorax group gene tonalli (tna) interacts genetically with the Brahma remodeling complex and encodes an SP-RING finger protein. Development 2003, 130:343–354.

51. Mohit P, Makhijani K, Madhavi MB, Bharathi V, Lal A, Sirdesai G, Reddy VR, Ramesh P, Kannan R, Dhawan J, Shashidhara LS: Modulation of AP and DV signaling pathways by the homeotic gene Ultrabithorax during haltere development in Drosophila. Dev Biol 2006, 291:356–367.

52. Beck S, Faradji F, Brock H, Peronnet F: Maintenance of Hox gene expression patterns. Adv Exp Med Biol 2010, 689:41–62.

53. Salvaing J, Lopez A, Boivin A, Deutsch JS, Peronnet F: The Drosophila Corto protein interacts with Polycomb-group proteins and the GAGA factor. Nucleic Acids Res 2003, 31:2873–2882.

54. Calgaro S, Boube M, Cribbs DL, Bourbon H-M: The Drosophila gene taranis encodes a novel trithorax group member potentially linked to the cell cycle regulatory apparatus. Genetics 2002, 160:547–560.

55. Lopez A, Higuet D, Rosset R, Deutsch J, Peronnet F: corto genetically interacts with Pc-G and trx-G genes and maintains the anterior boundary of Ultrabithorax expression in Drosophila larvae. Mol Genet Genomics 2001, 266:572–583.

56. Nègre N, Hennetin J, Sun LV, Lavrov S, Bellis M, White KP, Cavalli G: Chromosomal distribution of PcG proteins during Drosophila development. PLoS Biol 2006, 4:e170.

57. Ringrose L, Rehmsmeier M, Dura J-M, Paro R: Genome-Wide Prediction of Polycomb/Trithorax Response Elements in Drosophila melanogaster. Dev Cell 2003, 5:759–771.

58. Biggin MD, McGinnis W: Regulation of segmentation and segmental identity by Drosophila homeoproteins: the role of DNA binding in functional activity and specificity. Development 1997, 124:4425–4433.

59. Spitz F, Gonzalez F, Duboule D: A global control region defines a chromosomal regulatory landscape containing the HoxD cluster. Cell 2003, 113:405–417.

60. Grosveld F, van Assendelft GB, Greaves DR, Kollias G: Position-independent, high-level expression of the human beta-globin gene in transgenic mice. Cell 1987, 51:975–985.

61. Hnisz D, Abraham BJ, Lee TI, Lau A, Saint-André V, Sigova AA, Hoke HA, Young RA: Super-enhancers in the control of cell identity and disease. Cell 2013, 155:934–947.

62. Steffen PA, Ringrose L: What are memories made of? How Polycomb and Trithorax proteins mediate epigenetic memory. Nat Rev Mol Cell Biol 2014, 15:340–356.

63. Ringrose L, Paro R: Epigenetic regulation of cellular memory by the Polycomb and Trithorax group proteins. Annu Rev Genet 2004, 38:413–443.

64. Kwong C, Adryan B, Bell I, Meadows L, Russell S, Manak JR, White R: Stability and Dynamics of Polycomb Target Sites in Drosophila Development. PLoS Genet 2008, 4:e1000178.

65. Oktaba K, Gutiérrez L, Gagneur J, Girardot C, Sengupta AK, Furlong EEM, Müller J: Dynamic regulation by polycomb group protein complexes controls pattern formation and the cell cycle in Drosophila. Dev Cell 2008, 15:877–889.

66. de Boer E, Rodriguez P, Bonte E, Krijgsveld J, Katsantoni E, Heck A, Grosveld F, Strouboulis J: Efficient biotinylation and single-step purification of tagged transcription factors in mammalian cells and transgenic mice. Proc Natl Acad Sci USA 2003, 100:7480–7485.

67. Strübbe G, Popp C, Schmidt A, Pauli A, Ringrose L, Beisel C, Paro R: Polycomb purification by in vivo biotinylation tagging reveals cohesin and Trithorax group proteins as interaction partners. Proc Natl Acad Sci USA 2011, 108:5572–5577.

68. Garaulet DL, Foronda D, Calleja M, Sánchez-Herrero E: Polycomb-dependent Ultrabithorax Hox gene silencing induced by high Ultrabithorax levels in Drosophila. Development 2008, 135:3219–3228.

69. Petruk S, Sedkov Y, Riley KM, Hodgson J, Schweisguth F, Hirose S, Jaynes JB, Brock HW, Mazo A: Transcription of bxd noncoding RNAs promoted by trithorax represses Ubx in cis by transcriptional interference. Cell 2006, 127:1209–1221.

70. Schwartz YB, Kahn TG, Nix DA, Li X-Y, Bourgon R, Biggin M, Pirrotta V: Genome-wide analysis of Polycomb targets in Drosophila melanogaster. Nat Genet 2006, 38:700–705.

71. Bischof J, Maeda RK, Hediger M, Karch F, Basler K: An optimized transgenesis system for Drosophila using germ-line-specific phiC31 integrases. Proc Natl Acad Sci USA 2007, 104:3312–3317.

72. Bonn S, Zinzen RP, Perez-Gonzalez A, Riddell A, Gavin A-C, Furlong EE: Cell type-specific chromatin immunoprecipitation from multicellular complex samples using BiTS-ChIP. Nat Protoc 2012, 7:978–994.

73. Sandmann T, Jensen LJ, Jakobsen JS, Karzynski MM, Eichenlaub MP, Bork P, Furlong EEM: A Temporal Map of Transcription Factor Activity: Mef2 Directly Regulates Target Genes at All Stages of Muscle Development. Dev Cell 2006, 10:797–807.

74. Langmead B, Trapnell C, Pop M, Salzberg SL: Ultrafast and memory-efficient alignment of short DNA sequences to the human genome. Genome Biol 2009.

75. Kent WJ, Sugnet CW, Furey TS, Roskin KM, Pringle TH, Zahler AM, Haussler D: The human genome browser at UCSC. Genome Res 2002, 12:996–1006.

76. Tomancak P, Beaton A, Weiszmann R, Kwan E, Shu S, Lewis SE, Richards S, Ashburner M, Hartenstein V, Celniker SE, Rubin GM: Systematic determination of patterns of gene expression during Drosophila embryogenesis. Genome Biol 2002, 3:RESEARCH0088.

77. Schuettengruber B, Ganapathi M, Leblanc B, Portoso M, Jaschek R, Tolhuis B, van Lohuizen M, Tanay A, Cavalli G: Functional anatomy of polycomb and trithorax chromatin landscapes in Drosophila embryos. PLoS Biol 2009, 7:e13.

